# Voltage-sensitive sodium channel (*Vssc*) mutations associated with pyrethroid insecticide resistance in *Aedes aegypti* (L.) from Jeddah, Kingdom of Saudi Arabia – baseline information for a *Wolbachia* release program

**DOI:** 10.1101/2021.01.26.428347

**Authors:** Nancy M. Endersby-Harshman, AboElgasim Ali, Basim Alhumrani, Mohammed Abdullah Alkuriji, Mohammed B. Al-Fageeh, Abdulaziz Al-Malik, Mohammed S. Alsuabeyl, Samia Elfekih, Ary A. Hoffmann

## Abstract

**Background:** Dengue suppression often relies on control of the mosquito vector, *Aedes aegypti*, through applications of insecticides of which the pyrethroid group has played a dominant role. Insecticide resistance is prevalent in *Ae. aegypti* around the world and the resulting reduction of insecticide efficacy is likely to exacerbate the impact of dengue. Dengue has been a public health problem in Saudi Arabia, particularly in Jeddah, since its discovery there in the 1990s and insecticide use for vector control is widespread throughout the city. An alternative approach to insecticide use, based on blocking dengue transmission in mosquitoes by the endosymbiont *Wolbachia*, is being trialled in Jeddah following the success of this approach in Australia and Malaysia. Knowledge of insecticide resistance status of mosquito populations in Jeddah is a prerequisite for establishing a *Wolbachia*-based dengue control program as releases of *Wolbachia* mosquitoes succeed when resistance status of the release population is similar to that of the wild population.

**Methods:** WHO resistance bioassays of mosquitoes with deltamethrin, permethrin and DDT were used in conjunction with TaqMan^®^ SNP Genotyping Assays to characterise mutation profiles of *Ae. aegypti* from Jeddah.

**Results:** Screening of the voltage sensitive sodium channel (*Vssc*), the pyrethroid target-site, revealed mutations at codons 989, 1016 and 1534 in *Ae. aegypti* from two districts of Jeddah. The triple mutant homozygote (1016G/1534C/989P) was confirmed from Al Safa and Al Rawabi. Bioassays with pyrethroids (Type I and II) and DDT showed that mosquitoes were resistant to each of these compounds based on WHO definitions. An association between *Vssc* mutations and resistance was established for the Type II pyrethroid, deltamethrin, with one genotype (989P/1016G/1534F) conferring a survival advantage over two others (989S/1016V/1534C and the triple heterozygote). An indication of synergism of Type I pyrethroid activity with piperonyl butoxide suggests that detoxification by cytochrome P450s accounts for some of the pyrethroid resistance response in *Ae. aegypti* populations from Jeddah.

**Conclusions:** The results provide a baseline for monitoring and management of resistance as well as knowledge of *Vssc* genotype frequencies required in *Wolbachia* release populations to ensure homogeneity with the target field population.

## Background

Target-site resistance to pyrethroids in *Aedes aegypti*, also known as knockdown resistance (*kdr*), is an autosomal, incompletely recessive trait (1). *Vssc* mutations at codons 1016 and 1534 occur in *Ae. aegypti* within the pyrethroid receptor sites in Domains II (S6) and III (S6) of the protein molecule (2). A third mutation, S989P, which is often in perfect linkage with V1016G, is not known to reduce the sensitivity of the sodium channel (2), but confers some additive pyrethroid resistance in the homozygous state in combination with 1016G (3).

Pyrethroid resistance and *Vssc* mutations have been identified in *Ae. aegypti* from the Kingdom of Saudi Arabia and discussed in several studies (4–7). These three mutation sites are being screened routinely in samples of *Ae. aegypti* from Jeddah and are labelled as S989P, V1016G and F1534C (Figure 1) according to the codon numbering in the sequence of the most abundant splice variant of the house fly, *Musca domestica*, *Vssc* (GenBank accession nos. AAB47604 and AAB47605) (8). However, direct links between these mutations and resistance phenotypes in this locality are still unclear.

**Figure 1.**
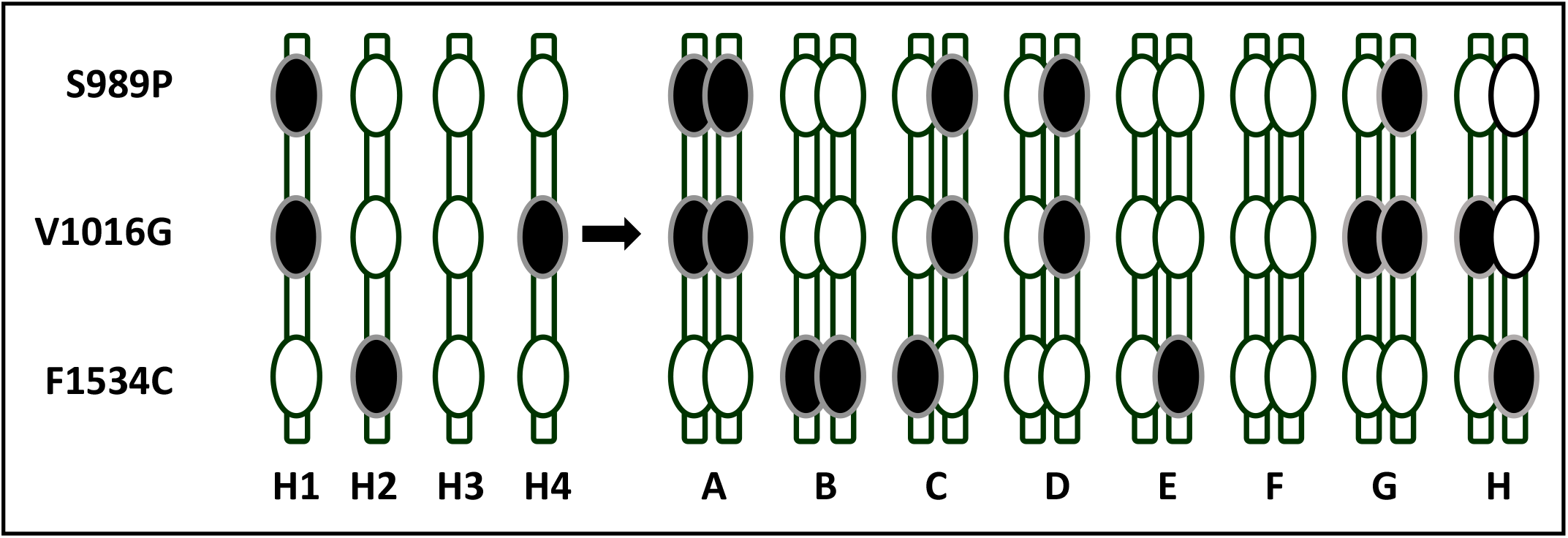
Indo-Pacific *Vssc* haplotypes and genotypes in *Aedes aegypti* (26)

Dengue has been a public health problem in Saudi Arabia and particularly in Jeddah since its discovery there in 1994 (9). Currently, dengue suppression relies on control of the mosquito vector, *Aedes aegypti*, through the applications of insecticides of which the pyrethroid group has played a dominant role (5). Insecticide resistance is prevalent in *Ae. aegypti* around the world (10, 11) and the resulting reduction of insecticide efficacy is likely to be exacerbating the impact of dengue and certainly threatens the long-term utility of this control method. Hence an alternative approach based on blocking dengue transmission in mosquitoes by the endosymbiont *Wolbachia* is being trialled following the success of this approach in suppressing dengue in other countries (12, 13). Preparation for a release of *Wolbachia*-infected mosquitoes for control of dengue transmission has commenced in Jeddah in the Kingdom of Saudi Arabia.

Knowledge of the insecticide resistance status of mosquito populations in Jeddah is one prerequisite for establishing a *Wolbachia*-based dengue control program. The presence and frequency of the mutants can be used as a measure of resistance in the local population of *Ae. aegypti* that can be compared with *Wolbachia*-infected mosquitoes destined for release. Mosquitoes are screened for mutations in the voltage-sensitive sodium channel (*Vssc*), the target site of pyrethroid insecticides and DDT (14) to characterise the population from the field for comparison with the future population for release. Similar releases of *Wolbachia* mosquitoes have been shown to succeed when insecticide resistance status of the release population is similar to that in the field (12, 15) and to fail or strike difficulties if the release population of mosquitoes does not carry equivalent resistance traits (16, 17). Although a *Wolbachia* program is not based on insecticide use, it is a reality that released mosquitoes will be exposed to insecticides in the field and therefore, must initially be able to survive in such an environment.

In this study we aimed:

1. To characterise the *Vssc* mutations 989/1016/1534 and haplotypes in the *Ae. aegypti* populations in Jeddah in two areas (Al Safa and Al Rawabi) to compare with known haplotype patterns in other parts of the world
2. To compare the distribution of these *Vssc* mutations in dead and surviving mosquitoes from bioassays with insecticides that target the *Vssc* (Type I and Type II pyrethroids as well as DDT which shows a similar mode of action to Type I pyrethroids (18))
3. To screen for the presence of a fourth mutation (codon 410) known to confer pyrethroid resistance to *Ae. aegypti* in a restricted number of locations (19–21) and, if found, to characterize this mutation in bioassay dead and surviving mosquitoes by sequencing a region of *Vssc* Domain I where this mutation may be located
4. To screen for a mutation in codon 1763, known from *Ae. aegypti* in Taiwan (22, 23) which does not affect the sensitivity of the *Vssc* (2), but may play a similar role to S989P in enhancing resistance conferred by the other mutations (22).
5. To use a synergist bioassay to find evidence of a metabolic component of pyrethroid resistance in *Ae. aegypti* from Jeddah

Knowledge about insecticide resistance in *Ae. aegypti* from the potential *Wolbachia* release sites is needed to ensure that similar traits are found in the strains of mosquitoes to be released to facilitate their survival under continued pressure of insecticide applications. This knowledge is also essential in developing long term insecticide resistance management strategies for Jeddah.

## Methods

### Sample collection

Mosquitoes were sampled in Al Safa-9 district located in central Jeddah and Al Rawabi district in the southern part of the city on three occasions (August 2018, October 2019 and February 2020) using oviposition bucket traps. The field sampling was performed over a two-week period for each operation and felts were collected twice and brought to the lab for rearing in separate containers, from which we tested an average of 10-12 mosquito samples (larval and adult stages). Mosquito colonies were built and maintained in the Jeddah lab using larvae collected from various collection points in Safa-9 district. Mosquitoes from these wild colonies were used for the WHO bioassays.

### WHO bioassays

The standard WHO insecticide resistance bioassay involving insecticide-impregnated papers was used for adult mosquitoes (24, 25). 25 female adults aged less than5 days post-eclosion and non blood fed were used as the test subjects. Five replicates and one control were used for each bioassay. Controls comprised the relevant solvent used for the insecticide papers (silicone oil for pyrethroids). The mosquitoes were exposed in WHO tubes to impregnated paper with diagnostic concentrations of deltamethrin (0.05%), permethrin (0.75%) or DDT (4%). Bioassays to test for synergism of permethrin 0.75% were conducted by exposing mosquitoes to piperonyl butoxide (PBO) (4%) for one hour followed by exposure to permethrin (0.75%) for an additional hour. Insecticide impregnated papers were purchased from the Vector Control Research Unit, Universiti Sains Malaysia, Penang. Knockdown was scored every ten minutes for a 1 h exposure period. After exposure, the mosquitoes were transferred into clean tubes and were supplied with cotton soaked in 10% sugar solution. Mortality was recorded after 24 h. Classification of resistance status was made based on pre-determined WHO guidelines in which mortality of <90% is characterized as resistant. Bioassays were repeated until at least 40 dead and 40 survivors could be collected for each insecticide to be used for analysis of *Vssc* mutations. Mosquitoes from the bioassays were stored in absolute ethanol.

### DNA extraction

DNA was extracted from adult mosquitoes or late instar larvae using either the DNeasy^®^ Blood and Tissue kit (QIAGEN Sciences, Maryland, USA), the Roche High Pure PCR template kit (Roche Molecular Systems, Inc., Pleasanton, CA, USA) according to the instructions of the manufacturer or extracted using Chelex^®^ 100 resin (Bio-Rad Laboratories Inc., Hercules, CA USA). Two final elutions of DNA were made from the kit extractions with the first being used for construction of genomic libraries for a related study and the second being used for screening of *Vssc* mutations after being diluted 1:5 with water. The same dilution factor was used for DNA extracted using Chelex^®^ 100 resin on samples which were not to be used for construction of genomic libraries.

### *Screening of* Vssc *mutations*

#### 989/1016/1534

Custom TaqMan^®^ SNP Genotyping Assays (Life Technologies, California, USA) were developed for each of the three target site mutations (codons 989, 1016, 1534) and were run on the Roche LightCycler^®^ 480 (26). Endpoint genotyping was conducted using the Roche LightCycler^®^ 480 Software Version 1.5.1.62.

#### 410 and 1763

A subset of samples was screened for a mutation at codon 410 in Domain I of the *Vssc* and codon 1763 in Domain IV by PCR and Sanger sequencing. Primers used to amplify the region around codon 410 (exon 10 in Domain I) of the *Vssc* were aegSCF10 (5’-GTGTTACGATCAGCTGGACC-3’ and aegSCR10 (5’-AAGCGCTTCTTCCTCGGC-3’) from Tancredi *et al*. (27). Primers to amplify codon 1763 in *Vssc* Domain IV were albSCF6 (5’-TCGAGAAGTACTTCGTGTCG-3’) and albSCR8 (5’-AACAGCAGGATCATGCTCTG-3’) (28). Amplicons were sequenced with albSCF7 (5’-AGGTATCCGAACGTTGCTGT-3’) by Macrogen Inc., South Korea for Sanger sequencing.

A 25 μL PCR master mix was set up as follows: 2.5 μL Standard Taq (Mg-free) Reaction Buffer (10x) (B9015 New England BioLabs Inc. Ipswich MA USA), 2.0 μL dNTP mix (10mM total) (Bioline (Aust) Pty Ltd, Eveleigh NSW Australia), MgCl_2_ (50 mM) (Bioline (Aust) Pty Ltd, Eveleigh NSW Australia), 1.25 μL of each primer (10 μM) (aegSCF10 5’-GTGTTACGATCAGCTGGACC-3’ and aegSCR10 5’-AAGCGCTTCTTCCTCGGC-3’ from), 0.125 μL IMMOLASE™ DNA Polymerase (5 u/μL) (Bioline (Aust) Pty Ltd, Eveleigh NSW Australia), 15.125 μL PCR-grade water and 2 μL DNA (1:10 dilution of Chelex^®^-extracted DNA). A PCR reaction of 95°C for 10 min, 35 cycles of 95°C 30 s, 55°C 45 s, 72°C 45 s and a final extension at 72°C for 5 min was run on an Eppendorf Mastercycler^®^ (Eppendorf AG, Hamburg Germany). Amplicons were observed on a 1.5% agarose gel (Bioline (Aust) Pty Ltd, Eveleigh NSW Australia) run for 40 min at 90 V stained with SYBR™ Safe DNA Gel Stain (Thermo Fisher Scientific Inc. Waltham MA USA) and viewed with a GelDoc XR Gel Documentation System (Bio-Rad Laboratories Inc. Hercules CA USA). Samples were sent to Macrogen Inc., South Korea, for Sanger sequencing. Amplicons for the codon 410 screen were sequenced with aegSCR10 and those for codon 1763 were sequenced with albSCF7 (5’-AGGTATCCGAACGTTGCTGT-3’). Sequences were analysed with Geneious Prime Version 2020 (Biomatters Ltd.).

Samples screened were those collected in Al Rawabi and Al Safa in February 2020. Sequences were first aligned to KY747529 (19) for verification as *Aedes aegypti Vssc* and then to KY747530.1 for codon 410 (mutant L) (19) and MK495874 (wildtype D) and MK495875 (mutant Y) (23) for codon 1763 for identification of a possible mutation.

### Statistical analyses

*Vssc* mutation data from Al Safa and Al Rawabi collected over three years were analyzed for site differences in genotype frequencies using contingency tables with significance tested through the chi-squared statistic. Monte Carlo tests were used to determine significance because expected values in some cells were particularly low even when data were combined across collection dates. These analyses were performed in IBM SPSS Statistics (IBM Corp., Armonk, NY, USA; 2013).

Odds ratios (with 95% confidence intervals) (29) were calculated to indicate the odds of an individual surviving exposure to an insecticide if it carried one particular genotype compared with another. An odds ratio of 1 indicates that there is no relationship between survival and the genotype under investigation. If 95% confidence intervals of the odds ratio do not span the value “1” then this suggests that the genotype is associated with survival.

## Results

### Vssc mutations in temporal field samples

An initial sample of ten individuals per district identified *Vssc* genotypes seen in samples from Asia and the Indo-Pacific except for one individual from Safa which did not fit the pattern (a homozygous mutant at codon 1016 and 989, but a heterozygote at 1534) (Table 1). Data from multiple studies (26, 30–33) suggest that certain haplotypes of the three mutation sites predominate in a population and there is little evidence of crossing over to disrupt the phase patterns found. The linkage patterns we have observed for these mutations in mosquitoes from Asia and the Indo-Pacific cannot produce individuals of genotype I found in Safa (Figure 1). To obtain this genotype, one parent had to contribute a triple mutant haplotype (H5) and the other contributed a second haplotype (H1) that we see in Asia/ Indo-Pacific samples (Figure 2). Saudi Arabia is one of the few countries where the triple mutant genotype has been reported previously (5) and it was found as a haplotype in the case here (two individuals of the same genotype) and also as four individuals made from a combination of the H5 and H2 haplotypes which we have called genotype J (Figure 2).

**Table 1.** Voltage-sensitive sodium channel (*Vssc*) mutations in *Aedes aegypti* from Al Safa and Al Rawabi districts, Jeddah, 2018, 2019, 2020

| Genotype |  | Safa |  | Safa |  | Safa |  | Rawabi |  | Rawabi |  | Rawabi |  |
| --- | --- | --- | --- | --- | --- | --- | --- | --- | --- | --- | --- | --- | --- |
|  |  | Aug-18 |  | Oct-19 |  | Feb-20 |  | Aug-18 |  | Oct-19 |  | Feb-20 |  |
|  |  | n | freq | n | freq | n | freq | n | freq | n | freq | n | freq |
| GG/GG/CC | K | 0 | 0.00 | 0 | 0.00 | 2 | 0.03 | 0 | 0.00 | 0 | 0.00 | 1 | 0.02 |
| GG/TG/CC | I | 1 | 0.10 | 1 | 0.02 | 2 | 0.03 | 0 | 0.00 | 0 | 0.00 | 0 | 0.00 |
| GG/TT/CC | A | 1 | 0.10 | 20 | 0.33 | 15 | 0.25 | 4 | 0.40 | 17 | 0.28 | 11 | 0.18 |
| TG/GG/TC | J | 0 | 0.00 | 0 | 0.00 | 2 | 0.03 | 0 | 0.00 | 0 | 0.00 | 1 | 0.02 |
| TG/TG/TC | C/L | 6 | 0.60 | 28 | 0.47 | 27 | 0.45 | 3 | 0.30 | 30 | 0.50 | 33 | 0.55 |
| TG/TT/TC | D | 0 | 0.00 | 0 | 0.00 | 0 | 0.00 | 0 | 0.00 | 0 | 0.00 | 1 | 0.02 |
| TT/TG/TT | E | 0 | 0.00 | 0 | 0.00 | 0 | 0.00 | 1 | 0.10 | 0 | 0.00 | 0 | 0.00 |
| TT/GG/TT | B | 2 | 0.20 | 11 | 0.18 | 12 | 0.20 | 2 | 0.20 | 13 | 0.22 | 13 | 0.22 |
| Total |  | 10 |  | 60 |  | 60 |  | 10 |  | 60 |  | 60 |  |

Test for site differences (all dates combined) $\chi^2=7.27$ , df=7, P=0.401 (0.388-0.413, 99% confidence intervals)

**Figure 2.**
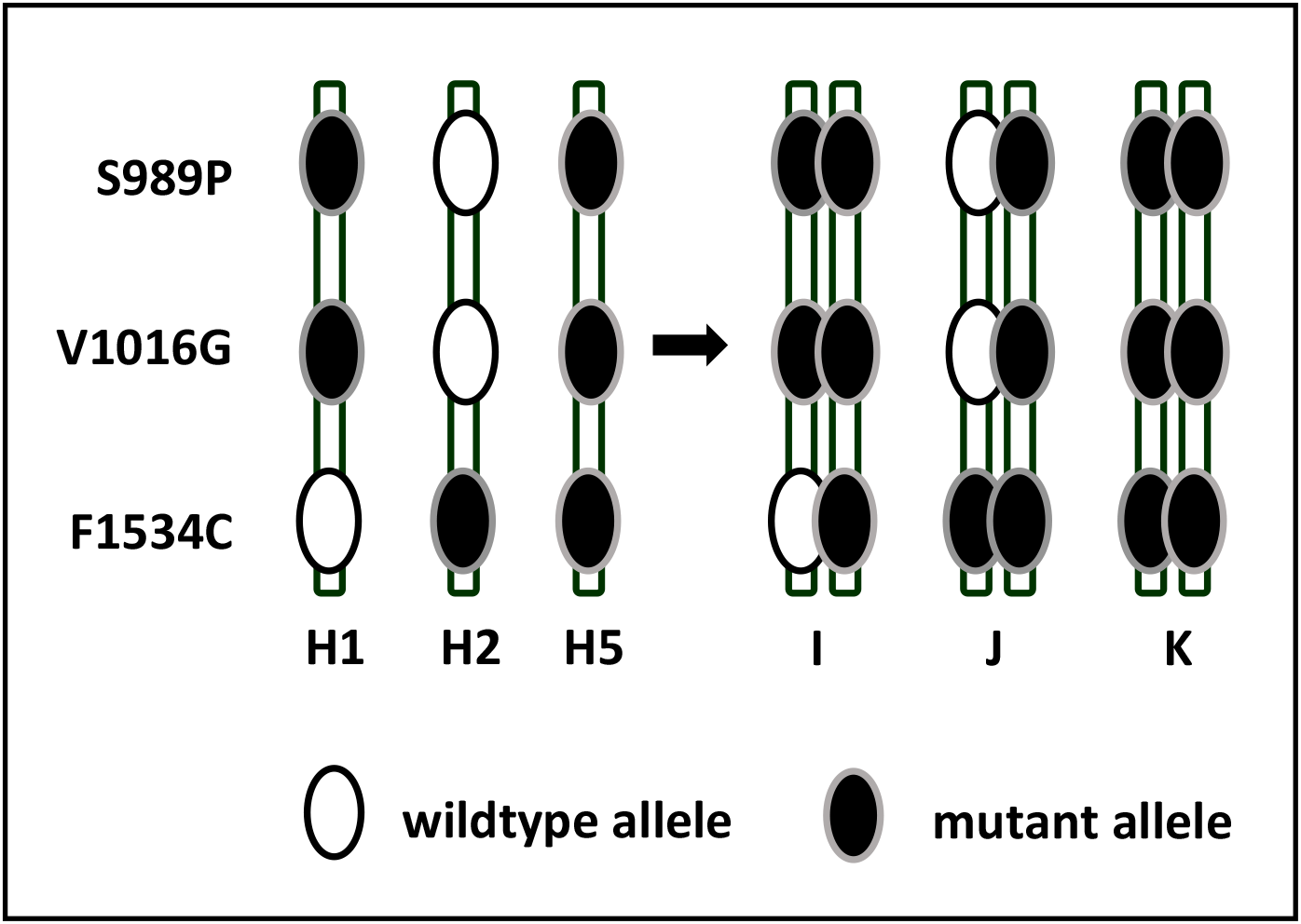
*Vssc* genotypes that include the triple mutant haplotype (H5) in *Aedes aegypti* from Saudi Arabia

The second screen consisted of sixty samples from each study site (Al Rawabi and Al Safa Districts, Jeddah), collected in October 2019. Three of the *Vssc* genotypes (A, B, C – Figure 1) found in the mosquitoes collected from these sites in Saudi Arabia were those found in samples from the Indo-Pacific region (26). However, there was again, one individual from Al Safa with genotype I (Table 1).

The third sample of *Ae. aegypti* was taken in February 2020 and again sixty samples from each site were screened for *Vssc* mutations. Genotype I was again found in mosquitoes from Al Safa (Table 1). Genotype J, which also contains the triple mutant haplotype (Figure 2), was found in both Al Safa and Al Rawabi and triple mutant homozygotes (genotype K – Figure 2) were found in both Al Safa (two individuals) and Al Rawabi (one individual). Two genotypes found only in the Al Rawabi sample (D and E – Figure 1 - one individual of each) indicate that the triple wildtype haplotype is present in that district, but was not found as a homozygote. The frequency of each of the common genotypes (A, B, C) is similar between Al Safa and Al Rawabi (Table 1), but there are differences in distribution of the rare genotypes (Figure 3). When data for all sampling dates were combined, there was no significant difference in genotypes between mosquitoes sampled in Al Safa and Al Rawabi (χ^2^=7.27, df=7, P=0.401, 0.388-0.413, 99% confidence intervals).

**Figure 3.**
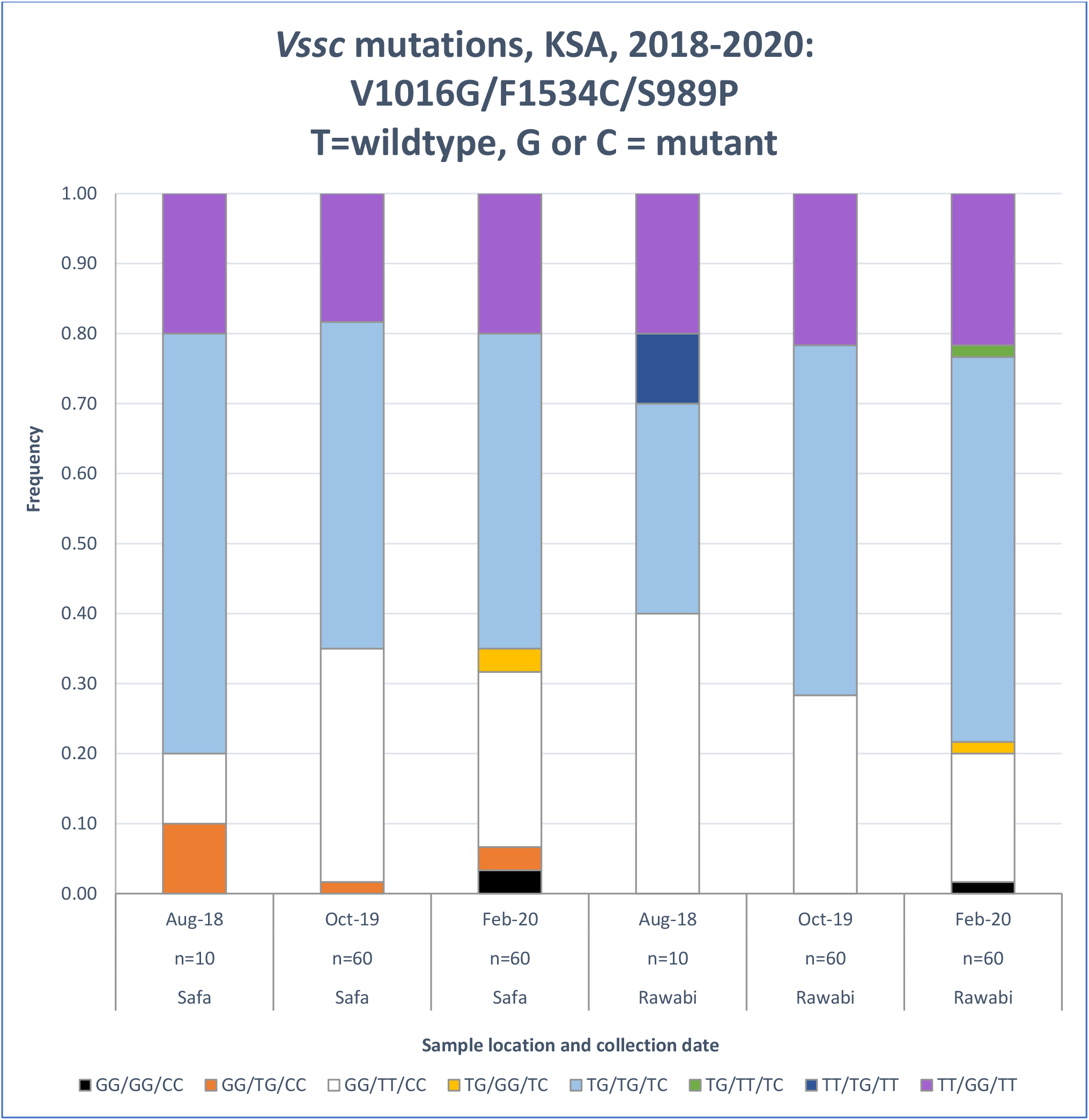
Comparison of frequency of *Vssc* genotypes in *Aedes aegypti* from Al Safa and Al Rawabi districts, Jeddah, Kingdom of Saudi Arabia

### WHO Bioassays

#### 1. Mortality

None of the bioassays conducted showed mortality in the range 98–100% which would indicate susceptibility according to WHO guidelines (34). Permethrin 0.75% induced only 44.8% mortality and DDT 4% bioassays showed mortality ranging from 3.2 to 10.4% confirming a high level of resistance to these compounds in *Ae. aegypti* from Jeddah. Four bioassays with deltamethrin 0.05% showed an average mortality just under 90% which indicates that the population of *Ae. aegypti* from Jeddah is resistant to this compound as well (Table 2).

**Table 2.** WHO bioassays of *Aedes aegypti* from Jeddah, KSA using permethrin 0.75%, deltamethrin 0.05%, DDT 4% - % mortality and numbers in *Vssc* screen

| Batch | Date | Status | Insecticide | Tube 1 | Tube 2 | Tube 3 | Tube 4 | Tube 5 | Total | Control | % mortality | Vssc screen |
| --- | --- | --- | --- | --- | --- | --- | --- | --- | --- | --- | --- | --- |
| 7 | 26-Sep-20 | DEAD | permethrin 0.75% | 10 | 8 | 17 | 13 | 8 | 56 | 0 | 44.8 | 40 |
| 7 | 26-Sep-20 | ALIVE | permethrin 0.75% | 15 | 17 | 8 | 12 | 17 | 69 | 25 |  | 40 |
| 1 | 2-Sep-20 | DEAD | deltamethrin 0.05% | 16 | 17 | 18 | 18 | 15 | 84 | 0 | 84.0 | 10 |
| 1 | 2-Sep-20 | ALIVE | deltamethrin 0.05% | 4 | 3 | 2 | 2 | 5 | 16 | 20 |  | 16 |
| 2 | 8-Sep-20 | DEAD | deltamethrin 0.05% | 20 | 18 | 18 | 18 | 19 | 93 | 1 | 92.5* | 10 |
| 2 | 8-Sep-20 | ALIVE | deltamethrin 0.05% | 0 | 2 | 2 | 2 | 1 | 7 | 19 |  | 7 |
| 4 | 15-Sep-20 | DEAD | deltamethrin 0.05% | 17 | 19 | 19 | 18 | 19 | 92 | 0 | 92.0 | 10 |
| 4 | 15-Sep-20 | ALIVE | deltamethrin 0.05% | 3 | 1 | 1 | 2 | 1 | 8 | 25 |  | 8 |
| 5 | 17-Sep-20 | DEAD | deltamethrin 0.05% | 19 | 18 | 17 | 18 | 18 | 90 | 0 | 90.0 | 10 |
| 5 | 17-Sep-20 | ALIVE | deltamethrin 0.05% | 1 | 2 | 3 | 2 | 2 | 10 | 20 |  | 10 |
| 6 | 26-Sep-20 | DEAD | DDT 4% | 2 | 2 | 3 | 2 | 4 | 13 | 0 | 10.4 | 8 |
| 6 | 26-Sep-20 | ALIVE | DDT 4% | 23 | 23 | 22 | 23 | 21 | 112 | 25 |  | 10 |
| 26 | 7-Nov-20 | DEAD | DDT 4% | 2 | 3 | 2 | 3 | 2 | 12 | 0 | 9.6 | 12 |
| 26 | 7-Nov-20 | ALIVE | DDT 4% | 23 | 22 | 23 | 22 | 23 | 113 | 25 |  | 10 |
| 28 | 8-Nov-20 | DEAD | DDT 4% | 1 | 1 | 0 | 1 | 2 | 5 | 0 | 4.0 | 7 |
| 28 | 8-Nov-20 | ALIVE | DDT 4% | 24 | 24 | 25 | 24 | 23 | 120 | 25 |  | 10 |
| 29 | 9-Nov-20 | DEAD | DDT 4% | 0 | 1 | 2 | 0 | 1 | 4 | 0 | 3.2 | 4 |
| 29 | 9-Nov-20 | ALIVE | DDT 4% | 25 | 24 | 23 | 24 | 24 | 120 | 25 |  | 10 |
| 31 | 12-Nov-20 | DEAD | DDT 4% | 1 | 2 | 1 | 0 | 1 | 5 | 0 | 4.0 | 6 |
| 31 | 12-Nov-20 | ALIVE | DDT 4% | 24 | 23 | 24 | 25 | 24 | 120 | 25 |  | 0 |
| 33 | 14-Nov-20 | DEAD | DDT 4% | 2 | 3 | 2 | 1 | 2 | 10 | 0 | 8.0 | 0 |
| 33 | 14-Nov-20 | ALIVE | DDT 4% | 23 | 22 | 23 | 24 | 23 | 115 | 25 |  | 0 |

#### 2. Synergist bioassay

Odds of mosquitoes being alive if exposed to permethrin 0.75% alone are just over twice those if exposed to permethrin 0.75% + PBO 4% (Table 3). PBO 4% alone caused some mortality, but the odds of being alive are not greater if exposed to permethrin compared with exposure to PBO alone (Table 3).

**Table 3.** WHO bioassays with permethrin 0.75% and synergist, piperonyl butoxide (PBO) and *Aedes aegypti* from Jeddah, KSA (OR = Odds Ratio, α=0.05, *=significance)

| Batch | Status | permethrin 0.75% | PBO 4% only | Insecticide + PBO 4% | Solvent control |
| --- | --- | --- | --- | --- | --- |
| 25 | Dead | 8 | 3 | 13 | 0 |
| 25 | Alive | 17 | 22 | 12 | 75 |
| 24 | Dead | 9 | 5 | 14 | 0 |
| 24 | Alive | 16 | 20 | 11 | 75 |
| 19 | Dead | 20 | 2 | 23 | 0 |
| 19 | Alive | 5 | 23 | 2 | 75 |
| **Total** | Dead | 37 | 10 | 50 | 0 |
|  | Alive | 38 | 65 | 25 | 225 |
| **% mortality** |  | 49.3 | 13.3 | 66.7 | 0.0 |
| | | **OR** | **Lower c.i.** | **Upper c.i.** | $\alpha=0.05$ |
| permethrin+PBO/permethrin |  | 0.49 | 0.25 | 0.94 |  |
| permethrin+PBO/PBO |  | 0.08 | 0.03 | 0.17 |  |
| permethrin/PBO |  | 0.16 | 0.07 | 0.35 |  |
| permethrin/permethrin+PBO |  | 2.05 | 1.06 | 3.97 | * |
| PBO/permethrin+PBO |  | 13.00 | 5.72 | 29.54 | * |
| PBO/permethrin |  | 6.33 | 2.83 | 14.16 | * |

#### 3. Comparison of *Vssc* genotypes in dead and survivors

Five *Vssc* genotypes (A, B, C/L, J, K) were identified in the surviving pool of mosquitoes screened and four genotypes (A, B, C/L, J) occurred in the dead mosquitoes exposed to 0.75% permethrin (Table 4). Two triple mutants were found in the survivors. There was no significant difference (α=0.05) in the odds of being alive after exposure to permethrin 0.75% if carrying one genotype over any other (Supplementary Table A).

**Table 4.**
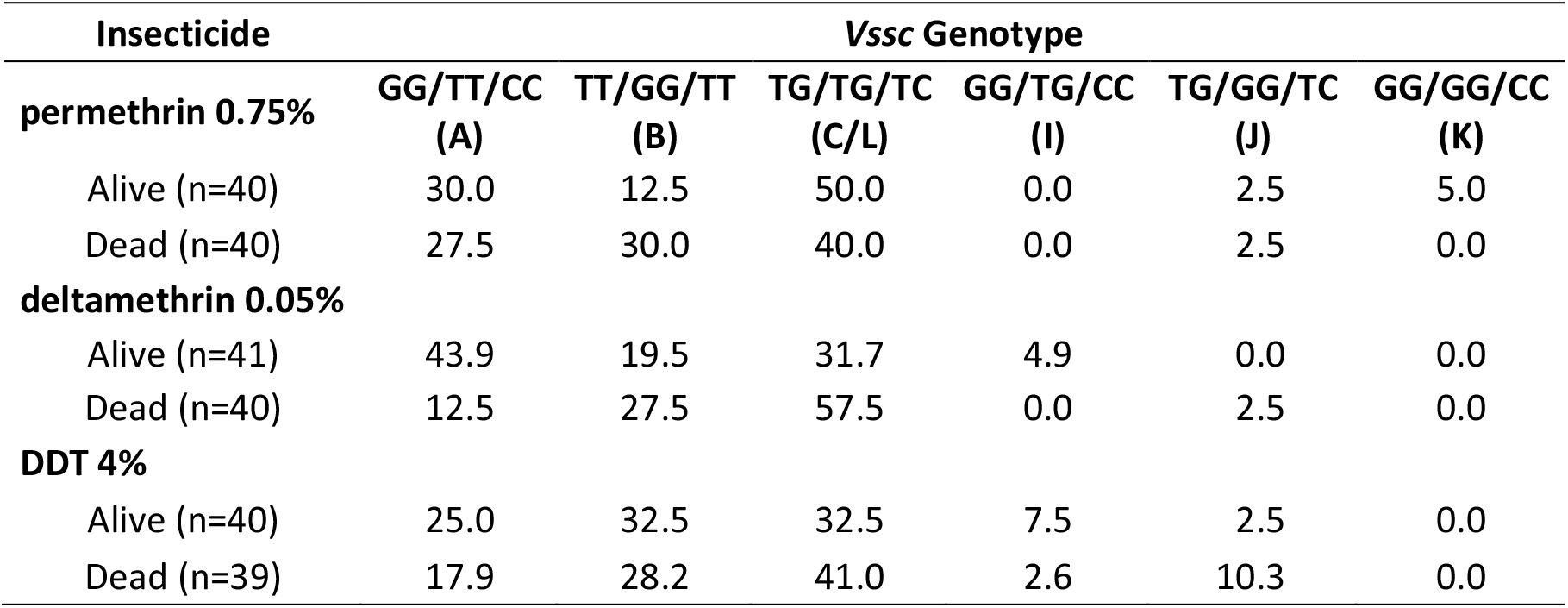
Frequency (%) of *Vssc* mutations in dead and surviving *Ae. aegypti* mosquitoes from Jeddah from WHO bioassays with permethrin (0.75%), deltamethrin (0.05%) or DDT (4%) (Order of mutations is 1016/1534/989, T=wildtype, G or C =mutant)

Survivors of deltamethrin 0.05% fell into four genotypes as did the dead individuals, though the fourth and least frequent genotype differed between them (I for alive and J for dead) (Table 4). Odds ratios were significant (α=0.05) for two genotype comparisons indicating that the odds of surviving were higher for individuals of genotype A than genotype B and also higher for genotype A than the triple heterozygote (genotype C or L) (Table 5).

**Table 5.** Odds of *Ae. aegypti* surviving with one *Vssc* genotype compared with another 24 h after a 1 h exposure to deltamethrin 0.05%. (OR =Odds Ratio with 95% confidence intervals) (*=significance, NS=not significant as confidence intervals encompass 1) (T=wildtype, G or C =mutant)

| **deltamethrin 0.05%** | **OR** | **LOWER** | **UPPER** | **$\alpha=0.05$** |
| --- | --- | --- | --- | --- |
| **GGTTCC/TTGGTT** | **4.95** | **1.29** | **19.01** | **\*** |
| **GGTTCC/TGTGTC** | **6.37** | **1.91** | **21.18** | **\*** |
| TTGGTT/TGTGTC | 1.29 | 0.41 | 4.01 | NS |
| TTGGTT/GGTTCC | 0.20 | 0.05 | 0.78 | NS |
| TGTGTC/GGTTCC | 0.16 | 0.05 | 0.52 | NS |
| TGTGTC/TTGGTT | 0.78 | 0.25 | 2.42 | NS |

Five genotypes (A, B, C/L, l, J) occurred in both survivors of DDT 4% and in the dead individuals (A, B, C/L, I, J) (Table 4). There was no significant difference (α=0.05) in the odds of being alive after exposure to DDT 4% if carrying one genotype over any other (Supplementary Table B).

### Codons 410 and 1763

32 sequences of 93-95 bp in length (30 from mosquitoes from Al Rawabi and two from Al Safa) were obtained for codon 410. Each sequence was wildtype at codon 410, coding for valine (V). 93 sequences of 117 bp in length were obtained for codon 1763 (47 from mosquitoes from Al Rawabi and 46 from Al Safa). Each sequence was identical and wildtype, coding for aspartic acid (D) at codon 1763.

## Discussion

*Vssc* mutations at codons 989, 1016 and 1534 are present in *Ae. aegypti* from two districts of Jeddah representing potential future release sites for *Wolbachia* mosquitoes for control of dengue. The populations are not fixed for one genotype, but genotypes A, B, C are common and no homozygote wildtype individuals were found in the sample. A wildtype haplotype exists, at least in Al Rawabi, so it is possible that homozygous wildtype individuals (genotype F) may occur in the population, albeit rarely. There is no indication that this haplotype still exists in Al Safa, but increased sampling might still find it. No instances of a mutation at codons 410 or 1763 were observed in the samples screened indicating lack of independent selection or contact of the mosquito populations with those from Taiwan (22), Brazil (19), Mexico (35) or West and Central Africa (20).

The triple mutant homozygote (1016G/1534C/989P) (genotype K – Figure 2) can now be confirmed from Al Safa and Al Rawabi, suggesting that *Ae. aegypti* from Saudi Arabia may have a unique population history which could be further explored in a full genomic analysis. The implications of finding the triple mutant as haplotype 5 and genotypes I – L are not well understood, but provide an opportunity to study effects of these genotypes in more detail in bioassays and potentially fitness experiments. Although the triple mutant genotype is rare in the populations of *Ae. aegypti* sampled from Jeddah (2-3%), the frequency is higher than that reported from Myanmar (0.98%) (32) and is high enough to facilitate collection of adequate numbers to isolate individuals for crossing experiments which could allow further characterisation of the H5 haplotype and I-L genotypes in bioassays. Two triple homozygous mutants were found in the live pool of our bioassay with permethrin 0.75%.

The *Vssc* genotypes found in Al Safa and Al Rawabi may all be constructed from haplotypes 1,2,3 and 5. Haplotype 4, found only in Taiwan of the countries sampled in the Indo-Pacific (26) and in Indonesia (3), is not found in the Saudi Arabian samples we have screened which is consistent with results of Al Nazawi, Aqili (5).

It is possible that the triple heterozygous genotype in Saudi Arabia could be composed of either H1 + H2 (usual condition in the Indo-Pacific – genotype C) or H3 + H5 (triple wildtype plus triple mutant – genotype L) (Figure 4). Both H3 and H5 are rare compared with H1 and H2, but there is a possibility that some of the heterozygotes have this rare configuration. It is not known whether there is any difference in insecticide resistance in mosquitoes which have one or the other haplotype combinations of the triple heterozygote.

**Figure 4.**
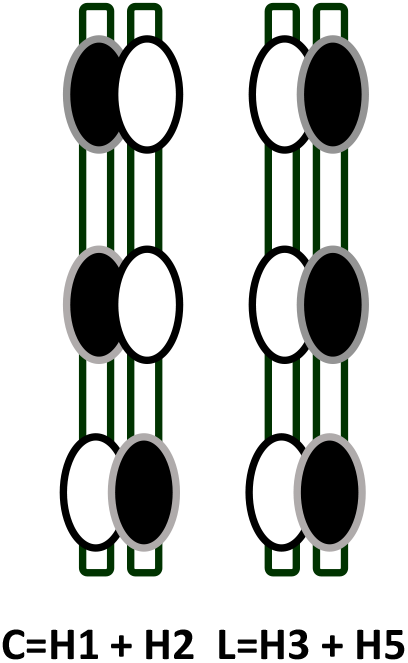
Putative configurations of *Vssc* triple mutant heterozygotes in *Aedes aegypti* from Saudi Arabia

Potential implications of the genotypes A-E and I-K on control of *Ae. aegypti* with pyrethroid insecticides have been surmised (Table 6) (5, 30, 36), but not all genotypes have been tested for effects on susceptibility. It is likely that efficacy of both Type I and Type II pyrethroids is compromised by many of these modifications to the *Vssc*, though genotypes have differential effects.

**Table 6.**
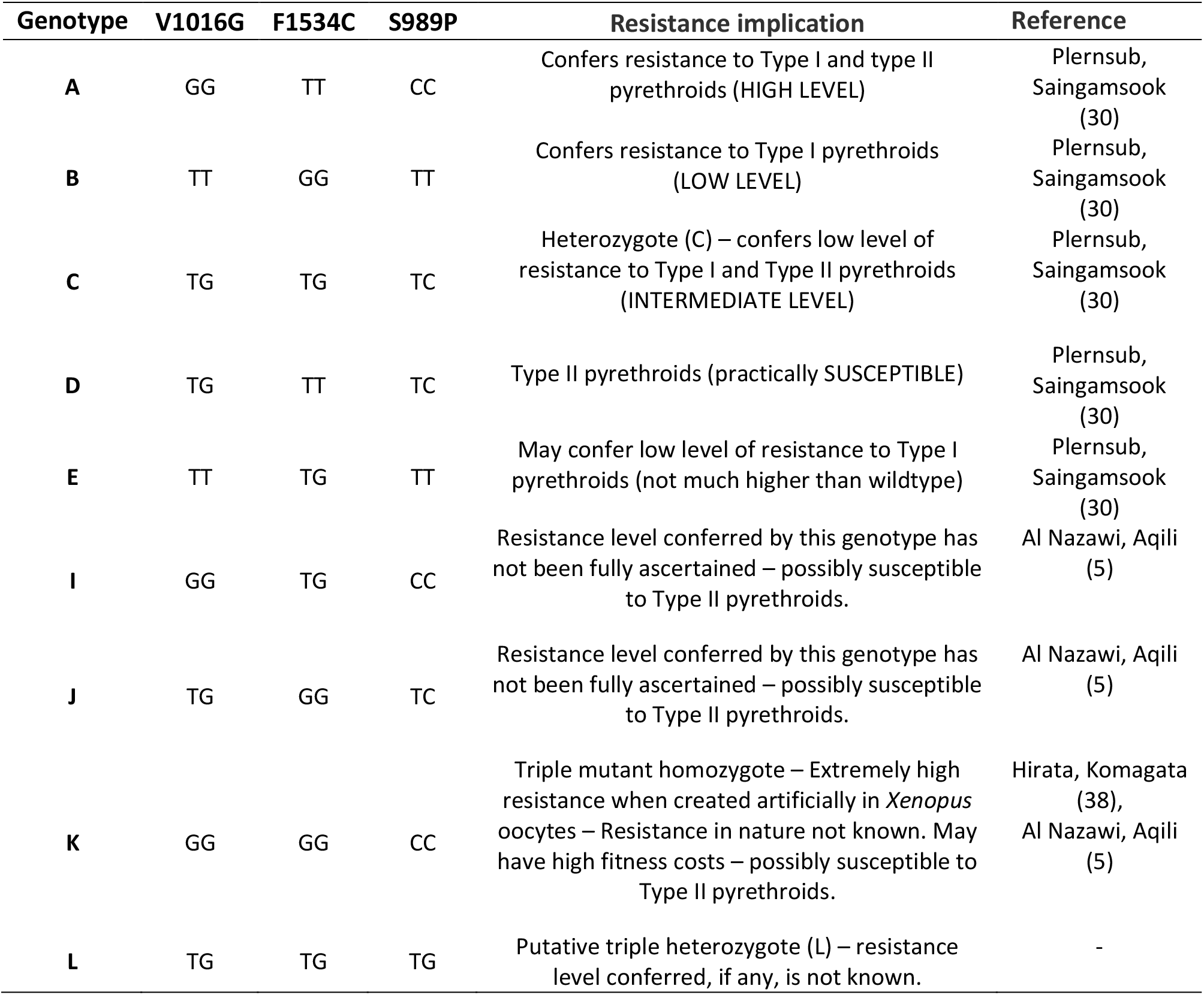
Pyrethroid resistance implications of voltage-sensitive sodium channel (*Vssc*) mutations in *Aedes aegypti* from the Kingdom of Saudi Arabia (Al Safa and Al Rawabi districts of Jeddah)

| Genotype | V1016G | F1534C | S989P | Resistance implication | Reference |
| --- | --- | --- | --- | --- | --- |
| **A** | GG | TT | CC | Confers resistance to Type I and type II pyrethroids (HIGH LEVEL) | Plernsub, Saingamsook (30) |
| **B** | TT | GG | TT | Confers resistance to Type I pyrethroids (LOW LEVEL) | Plernsub, Saingamsook (30) |
| **C** | TG | TG | TC | Heterozygote (C) – confers low level of resistance to Type I and Type II pyrethroids (INTERMEDIATE LEVEL) | Plernsub, Saingamsook (30) |
| **D** | TG | TT | TC | Type II pyrethroids (practically SUSCEPTIBLE) | Plernsub, Saingamsook (30) |
| **E** | TT | TG | TT | May confer low level of resistance to Type I pyrethroids (not much higher than wildtype) | Plernsub, Saingamsook (30) |
| **I** | GG | TG | CC | Resistance level conferred by this genotype has not been fully ascertained – possibly susceptible to Type II pyrethroids. | Al Nazawi, Aqili (5) |
| **J** | TG | GG | TC | Resistance level conferred by this genotype has not been fully ascertained – possibly susceptible to Type II pyrethroids. | Al Nazawi, Aqili (5) |
| **K** | GG | GG | CC | Triple mutant homozygote – Extremely high resistance when created artificially in *Xenopus* oocytes – Resistance in nature not known. May have high fitness costs – possibly susceptible to Type II pyrethroids. | Hirata, Komagata (38), Al Nazawi, Aqili (5) |
| **L** | TG | TG | TG | Putative triple heterozygote (L) – resistance level conferred, if any, is not known. | - |

Although resistance to three insecticides which target the *Vssc* was detected in bioassays in the *Ae. aegypti* from Jeddah in this study, the only clear association between *Vssc* mutations and resistance was established for the Type II pyrethroid, deltamethrin, with genotype A conferring a survival advantage compared with genotypes B (1534C) or C (triple heterozygote). Genotype A is known to reduce the sensitivity of the *Vssc* for both Type I and Type II pyrethroids (2), but in our study there was no obvious effects for Type I (permethrin 0.75%) or DDT. It may be that this lack of effect is concentration related and a different concentration of permethrin or DDT would reveal a segregation of genotypes between dead and surviving mosquitoes as reported from the Jazan region of Saudi Arabia (4). In a similar study in *Ae. aegypti* from Malaysia (15), a survival advantage over wildtype was conferred to mosquitoes by genotypes B and C when exposed to permethrin 0.25%.

It is important to note that target-site resistance may not be the only mechanism of resistance to pyrethroids that has been selected in the Jeddah populations of *Ae. aegypti*. A role of metabolic resistance in *Ae. aegypti* from Al Safa and Al Rawabi is likely. Al Nazawi et al. (5) saw an increase in mortality of *Ae. aegypti* from Jeddah with deltamethrin, using the synergist (PBO), which indicates that detoxification by oxidases (cytochrome P450 mixed function oxidase system) accounts for some of the pyrethroid resistance response. We also found an indication of such synergism in bioassays with the Type I pyrethroid, permethrin 0.75%.

## Conclusions

The continuing presence of the wildtype haplotype and minor difference in genotypes between Al Safa and Al Rawabi suggests that selection for resistance may be patchy between districts. Changes in genotype frequencies over time also suggest that selection for pyrethroid resistance is ongoing. The results provide a baseline for ongoing monitoring of resistance particularly with implementation of resistance management programs and also indicate *Vssc* genotypes required in *Wolbachia* release populations to ensure homogeneity with the target field population.

## Declarations

### Ethics approval and consent to participate

not applicable

### Consent for publication

not applicable

### Availability of data and materials

All data generated or analysed during this study are included in this published article [and its supplementary information files].

### Authors’ contributions

NMEH, AAH and SE designed the study and wrote the paper. AEA conducted all WHO bioassays. NMEH conducted DNA extraction, SNP genotyping and sample preparation for DNA sequencing. NMEH and AAH analysed the data. BB, MA, MAF, AAM and MSA provided research services for the study and reviewed the manuscript.

## Acknowledgements

Thanks to the field operations teams in Safa and Rawabi districts, led by Dr Abdulghaffar Azhari, for collecting mosquitoes and to all team members involved in DNA extractions and bioassays

