## Supplemental Tables for "Voltage-sensitive sodium channel (*Vssc*) mutations associated with pyrethroid insecticide resistance in *Aedes aegypti* (L.) from Jeddah, Kingdom of Saudi Arabia – baseline information for a *Wolbachia* release program"

**Additional File 1**

Supplementary Table A. Frequency of *Vssc* mutations in dead and surviving *Ae. aegypti* mosquitoes from WHO insecticide paper bioassays with Type I pyrethroid, permethrin (0.75%) (OR =Odds Ratio with 95% confidence intervals)

| | **GG/TT/CC** | **TT/GG/TT** | **TG/TG/TC** | **GG/TG/CC** | **TG/GG/TC** | **GG/GG/CC** | |
|---|---|---|---|---|---|---|---|
| Alive | 12 | 5 | 20 | 0 | 1 | 2 | 40 |
| Dead | 11 | 12 | 16 | 0 | 1 | 0 | 40 |

| DDT 4% | **OR** | **LOWER** | **UPPER** | $\alpha$=0.05 |
|---|---|---|---|---|
| GGTTCC/TTGGTT | 2.62 | 0.70 | 9.86 | NS |
| GGTTCC/TGTGTC | 0.87 | 0.31 | 2.49 | NS |
| TTGGTT/TGTGTC | 0.33 | 0.10 | 1.14 | NS |
| TTGGTT/GGTTCC | 0.38 | 0.10 | 1.44 | NS |
| TGTGTC/GGTTCC | 1.15 | 0.40 | 3.27 | NS |
| TGTGTC/TTGGTT | 3.00 | 0.87 | 10.30 | NS |

Supplementary Table B. Frequency of *Vssc* mutations in dead and surviving *Ae. aegypti* mosquitoes from WHO insecticide paper bioassays with DDT (4%) (OR =Odds Ratio with 95% confidence intervals)

| | **GG/TT/CC** | **TT/GG/TT** | **TG/TG/TC** | **GG/TG/CC** | **TG/GG/TC** | **00/TT/00** | **TOTAL** |
|---|---|---|---|---|---|---|---|
| Alive | 10 | 13 | 13 | 3 | 1 | 0 | 40 |
| Dead | 7 | 11 | 16 | 1 | 4 | 1 | 40 |

| DDT 4% | **OR** | **LOWER** | **UPPER** | $\alpha$=0.05 |
|---|---|---|---|---|
| GGTTCC/TTGGTT | 1.21 | 0.34 | 4.24 | NS |
| GGTTCC/TGTGTC | 1.76 | 0.52 | 5.91 | NS |
| TTGGTT/TGTGTC | 1.45 | 0.49 | 4.31 | NS |
| TTGGTT/GGTTCC | 0.83 | 0.24 | 2.91 | NS |
| TGTGTC/GGTTCC | 0.57 | 0.17 | 1.91 | NS |
| TGTGTC/TTGGTT | 0.69 | 0.23 | 2.04 | NS |
| GGTTCC/GGTGCC | 0.48 | 0.04 | 5.58 | NS |
| GGTTCC/TGGGTC | 5.71 | 0.52 | 62.66 | NS |
| TTGGTT/GGTGCC | 0.39 | 0.04 | 4.35 | NS |
| TTGGTT/TGGGTC | 4.73 | 0.46 | 48.77 | NS |
| GGTGTC/GGTTCC | 2.10 | 0.18 | 24.60 | NS |
| GGTGTC/TTGGTT | 2.54 | 0.23 | 28.02 | NS |
| GGTGTC/TGTGTC | 3.69 | 0.34 | 39.84 | NS |
| GGTGTC/TGGGTC | 12.00 | 0.51 | 280.11 | NS |
| TGGGTC/GGTTCC | 0.18 | 0.02 | 1.92 | NS |
| TGGGTC/TTGGTT | 0.21 | 0.02 | 2.18 | NS |
| TGGGTC/TGTGTC | 0.31 | 0.03 | 3.10 | NS |
| TGGGTC/GGTGCC | 0.08 | 0.00 | 1.95 | NS |
